# Phylogeny of cortical-hippocampal projections reveals selective elimination of input from sensory regions

**DOI:** 10.1101/2024.05.09.593130

**Authors:** Daniel Reznik, Piotr Majka, Marcello G P Rosa, Menno P Witter, Christian F Doeller

## Abstract

The study of cross-species differences in the organization of the cerebral cortex received significant attention during the past century. However, the evolutionary principles governing changes in brain connectivity and their impact on cognition remain poorly understood. In the current study, we analysed differences in anatomical input from the cerebral cortex to the hippocampal region across six species spanning more than 100 million years of mammalian evolution - the tenrec, rat, cat, marmoset, macaque, and human. Our results demonstrate that unlike direct transmodal cortical input, direct unimodal cortical input to the hippocampal region was selectively eliminated with the increase in brain size. Additionally, input from primary sensory regions was eliminated prior to input from non-primary sensory regions. Our findings suggest that hippocampus-related processes in different species operate on fundamentally different types of information, potentially underpinning cross-species differences in hippocampus-dependent cognition. Furthermore, our findings outline the evolutionary trajectory of the anatomical circuitry associating the human hippocampal region with the neocortex.

## Introduction

Cross-species differences in cognition are typically attributed to the changes that occurred to the cerebral cortex through evolution (Krubitzer, 2007, 2009; Reardon et al., 2018). This includes changes in size, structural differentiation and functional specialization of the cortex into increasingly larger number of cortical areas, which have been hypothesised to enable an increase in the animal’s behavioral range, its flexibility and complexity (Kaas, 1989). Comparative research into the mammalian lineage has demonstrated that even though evolution of the cerebral cortex followed an ancient phylogenetic template, the relative size of cortical areas involved in non-sensory processing dramatically increased through mammalian evolution, especially in humans, playing a major role in the development of unprecedented high-order cognitive functions (Buckner, 2026; Buckner & Krienen, 2013; Krubitzer, 2009; Smaers et al., 2021). Over the past century, significant attention has been devoted to the study of differences in the organization of the cortex across species, and to the potential contribution of this factor to cross-species differences in cognition (Brodmann, 1909; Buckner & Krienen, 2013; Chaplin et al., 2013; Kaas, 1989; Krubitzer, 2007; Orban et al., 2004; Petersen et al., 2024; M. Rosa & Manger, 2005; Smaers et al., 2021). However, the notion that animal behavior is the product of coordinated activity between anatomically interconnected brain regions (Bargmann & Marder, 2013; Goldman-Rakic, 1988) was rarely addressed from a comparative perspective. Hence, the evolutionary principles associating changes in anatomical connectivity with cross-species variations in cognition remain poorly understood.

Anatomical studies in mammals indicate that whole-brain cortical connectivity converges into the hippocampal region, comprising the hippocampal formation, perirhinal cortex, entorhinal cortex and parahippocampal cortex (postrhinal in the rodent) (Felleman & Van Essen, 1991; Witter et al., 1989). Consequently, the hippocampal region is critically implicated in cognitive functions requiring large-scale multimodal integration, such as memory, navigation, future planning and social cognition (Allen & Fortin, 2013; Eichenbaum, 2000, 2017; Manns & Eichenbaum, 2006). The complex nature of intrinsic anatomical organization and connections of the hippocampal region provided multiple opportunities for phenotypical divergence across mammals. However, in striking contrast to the cerebral cortex, the gross anatomical, cytoarchitectural and functional properties of the hippocampal region have remained largely conserved during 200 million years of mammalian evolution (Amaral & Witter, 1989; Manns & Eichenbaum, 2006; Witter et al., 1989). This remarkable preservation of the hippocampal region across species is consistent with its role as a major connectivity anchor positioned at the pinnacle of the distributed cortical hierarchy (Burwell, 2000; Felleman & Van Essen, 1991; Reznik et al., 2024). These anatomical and functional properties of the hippocampal region raise the possibility that it performs similar intrinsic computations across species, albeit operating on different types of cortical input (Kaas, 2019; Manns & Eichenbaum, 2006), which could potentially contribute to cross-species differences in hippocampus-dependent functions (Jeffery et al., 2024; Martinez-Trujillo, 2025; Rolls, 1999). This notion received preliminary support from comparing cortical connectivity of the hippocampal region across rats and macaques (Burwell et al., 1995), and across macaques and humans (Eichert et al., 2024; Reznik et al., 2023), suggesting that some connectivity patterns are preserved, while others have changed across species (Assaf et al., 2020; Herculano-Houzel et al., 2010; Ringo, 1991; Suárez et al., 2018).

In the current study, we leveraged the contrast between the preserved hippocampal region and the diverse cortex to test whether the type of information funnelled into the hippocampal region systematically changed through evolution. To this end, we examined differences in cortical input to the hippocampal region from unimodal and transmodal cortical areas across all mammalian species with known connectivity of the hippocampal region – the tenrec (*Echinops telfairi),* rat *(Rattus norvegicus),* cat *(Felis catus),* marmoset *(Callithrix jacchus),* macaque *(Macaca fascicularis),* and human *(Homo sapiens)*. These six species represent branches of the mammalian evolutionary tree that diverged over more than 100 million years of evolution (Figure 1). We analyzed anatomical tract-tracing data for the tenrec, rat, cat, marmoset and macaque, and functional magnetic resonance imaging (fMRI) data for humans, as a proxy of anatomical connectivity (Liu et al., 2019; Vincent et al., 2007, 2010). Our analyses point to gradual elimination of direct anatomical input from unimodal sensory regions and selective preservation of direct input from transmodal cortical regions, thus potentially outlining a major trend in the evolution of cortico-hippocampal circuitry during mammalian phylogeny.

**Figure 1.**
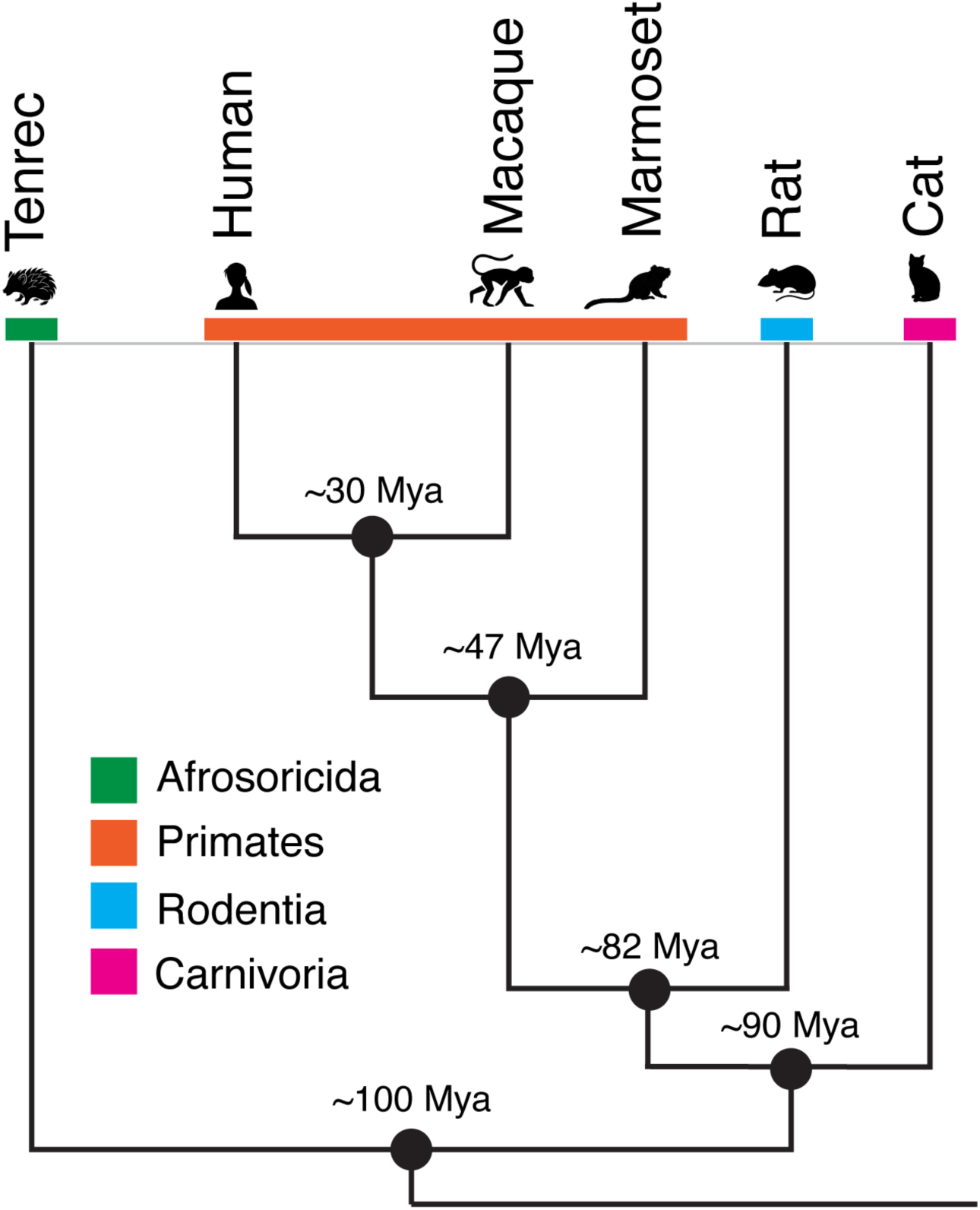
Mammalian species used in this study. We analyzed tract-tracing data from the tenrec, rat, cat, marmoset and macaque, which cover around 100 million years of mammalian evolution. Note that despite long periods of independent evolution, tenrecs and rats have brains with relatively similar sizes, whereas the marmoset split from the common ancestor with the macaque relatively recently, but have a much smaller and smoother brain compared with macaques (Heuer et al., 2025).

## Materials and Methods

The central aim of this study was to provide a cross-species comparative framework to the changes in anatomical connectivity associating the mammalian cortex with the hippocampal region. Due to the dramatic differences in the structural properties of the cortex across species, it is a challenging task to define homologous cortical regions that are present in all species, especially when cortical regions involved in non-sensory processing are considered (Brodmann, 1909; M. Rosa & Manger, 2005; Uylings et al., 2003). Therefore, we avoided comparing input from potential cortical homologues across species; instead, we utilised the phylogenetically conserved property of cortical hierarchy and divided the cortex of each species into cortical regions involved in unimodal sensory processing and cortical regions involved in transmodal processing. Unimodal cortical areas were defined as areas that were shown to selectively respond predominantly to one type of sensory modality, which include piriform, visual, auditory and somatosensory cortices. Where possible, unimodal sensory cortical regions were divided into primary sensory regions and non-primary sensory regions. Primary sensory regions, with the exception of the piriform cortex, were defined by direct input from sensory thalamic nuclei; non-primary sensory regions were defined by lack of connections with primary thalamic nuclei and by input from primary sensory regions. Transmodal cortical regions were defined as regions that do not to show specificity to any single sensory modality, which included polymodal and limbic cortical regions (Mesulam, 1998). See Supporting Information for more details about the mapping of unimodal sensory and transmodal regions in each species.

In the tenrec, we used data from studies that examined anatomical connections of the tenrec hippocampal region (Künzle, 2002, 2003). In the rat, we used a prior analysis of more than 16,000 connectivity reports, including the hippocampal region (Bota et al., 2015), as well as data from a study that examined input to the hippocampal region using retrograde tracer injections (Burwell & Amaral, 1998). In the marmoset we used an online brain connectivity atlas, which summarizes data from multiple fluorescent retrograde tracer injections (www.marmosetbrain.org; (Majka et al., 2016, 2020). Additionally, we used connectivity data that are not yet publicly available and will be added to the marmoset online atlas in future releases. In the cat, similar to the rat, we used a previous analysis of cortical connectivity, including the hippocampal region, derived from more than 90 anatomical connectivity studies (Scannell et al., 1995), as well as data from a study that examined connectivity between the cat’s prepiriform cortex and entorhinal cortex (Krettek & Price, 1977). In the macaque, we used studies that examined input to the hippocampal region using retrograde tracer injections, as well as studies that examined output projections of distributed cortical regions, including to the hippocampal region, using anterograde tracer injections (Cavada & Goldman-Rakic, 1989; Ding et al., 2000; Insausti et al., 1987; Lavenex et al., 2002; Mohedano-Moriano et al., 2007; Muñoz & Insausti, 2005; Seltzer & Pandya, 1984; Suzuki & Amaral, 1994). In humans, we used recent precision fMRI connectivity studies that examined the functional architecture of the hippocampal region at individual-subject level (Angeli et al., 2025; Edmonds et al., 2024; Reznik et al., 2023, 2024; Zheng et al., 2021).

First, in each species, we calculated the proportion of the cortical mantle dedicated to unimodal sensory processing and to transmodal processing (table S1-S6). To this end, previous cortical parcellations were used for each species. Next, using existing tract-tracing (for non-human animals) and fMRI connectivity (for humans) data, we calculated the overall proportion of the cortex, collapsed across unimodal and transmodal regions, projecting to the hippocampal region (or associated with it, in case of fMRI data). Connectivity was considered for each separate cortical region in each species. Since tracer injections or labelled cells often target a portion of different subregions, the entire subregion was considered to have connections with the hippocampal region even if only a portion was examined or showed labelled cells.

Lastly, we calculated separately the proportion of unimodal and transmodal regions projecting to the hippocampal region in each species. Importantly, whereas in the tenrec, cat, rat and macaque the anatomical projections to the hippocampal region were estimated directly by examining labelling of cells following tracer injections into the hippocampal region or distributed cortical regions, in the marmoset and human, cortical projections to the hippocampal region were estimated indirectly (see Supporting Information for more details).

To account for methodological differences in quantification of animal tracing data, we included all anatomical connections, including weak cortical input to the hippocampal region (e.g., 1-5% connectivity in the macaque; connections that were originally described as occasional or sporadic were not included in the analysis). In the tenrec, since the only studies that reported anatomical connectivity of the hippocampal region used a subjective rating scale (Künzle, 2002, 2003), only connections that were rated 3 (moderate connectivity) or above were included in the analysis. In the marmoset, since only data on anatomical projections from the hippocampal region are available (Majka et al., 2020), anatomical projections to the hippocampal region were estimated based on the reciprocity of the anatomical connections between the hippocampal region and the broader neocortex in the macaque (Lavenex et al., 2002). To account for weaker return projections from the hippocampal region to the primate cortex (Lavenex et al., 2002), we used a connectivity threshold of log_10_(FLNe) = -3 (fraction of extrinsic labelled neurons; average value across all available experiments in the same cortical area) in the marmoset. Only connections that showed up in more than half of the experiments for each cortical area in the marmoset were considered. For calculating the final estimate of marmoset projections from the neocortex to the hippocampal region, we averaged the percentage of input projections estimated solely on output projections and the percentage of input projections estimated using anatomical data in the macaque (see Supporting Information for more details, including Figure S1 for analysis without macaque-based adjustment of marmoset data).

For interpreting the associations of the human hippocampal region with the cerebral cortex estimated with fMRI connectivity we used a network-based approach, meaning that only distributed networks that were previously reported to associate with the human hippocampal region were included in the analysis (but see (Bergmann et al., 2016) and (S.-F. Wang et al., 2016)). It is important to emphasize that human connectivity data should be interpreted in light of our methodological choice to focus on fMRI connectivity findings. Other non-invasive methods, such as diffusion-weighted imaging (DWI), provide as well important insights into the anatomical organization of the human hippocampal region (Dalton et al., 2022; Huang et al., 2021; Maller et al., 2019). Specifically, while fMRI connectivity did not reveal associations between the human hippocampal region and early visual areas (Reznik et al., 2023), a recent DWI connectivity study points to such visual-hippocampal associations (Dalton et al., 2022).

Following recent findings demonstrating that animals with similar brain sizes show similar connectivity and behavioral phenotypes, irrespective of their phylogenetic similarity (Heuer et al., 2025), we examined how changes in cortical input to the hippocampal region are associated with species’ cortical surface area (log-transformed), without taking phylogenetic relationships into account. We used previous studies to estimate cortical surface area in the tenrec (Krubitzer et al., 1997), rat (Riddle et al., 1992), marmoset (Majka et al., 2016), cat (Rathjen et al., 2003), macaque (Felleman & Van Essen, 1991) and human (Donahue et al., 2018).

## Results

We examined cross-species changes in anatomical projections from the cortex to the hippocampal region in six different species (Figure 1). Our results indicate that the percentage of total cortical area projecting to the hippocampal region reduces with the increase in cortical surface area (Figure 2a; Pearson correlation coefficient = -0.94, p<0.01). In small non-primate mammals – the tenrec and rat – nearly the entire cortical mantle directly projects to the hippocampal region. However, in the marmoset, macaque and human, cortical subregions corresponding to about half of the cortex project to the hippocampal region. Next, we calculated direct input to the hippocampal region from unimodal and transmodal cortical regions relative to the total unimodal and transmodal cortical area in each species. We observed that relative input from each modality reduced with the increase in cortical surface area. Critically, this reduction disproportionally targeted input from unimodal cortical areas, while the input from transmodal areas was relatively preserved across species (Figure 2b; repeated measures ANCOVA, F_(1,4)_=22.84 p<0.01; see Figure S2 for unimodal and transmodal input of total cortical surface; see Figure S3 for an analysis using only tract tracing data, excluding humans). For example, in the rat, all primary and non-primary sensory regions have monosynaptic input to the hippocampal region. On the other hand, in larger mammals, such as cat and primates, we observed a gradual reduction in direct cortical input from unimodal sensory regions. Importantly, these changes in connectivity could not be explained merely by changes in the relative size of transmodal areas in different species. Even though about 1/3 of the human brain is involved in unimodal sensory processing, only the piriform cortex might be associated with the hippocampal region (see Discussion).

**Figure 2.**
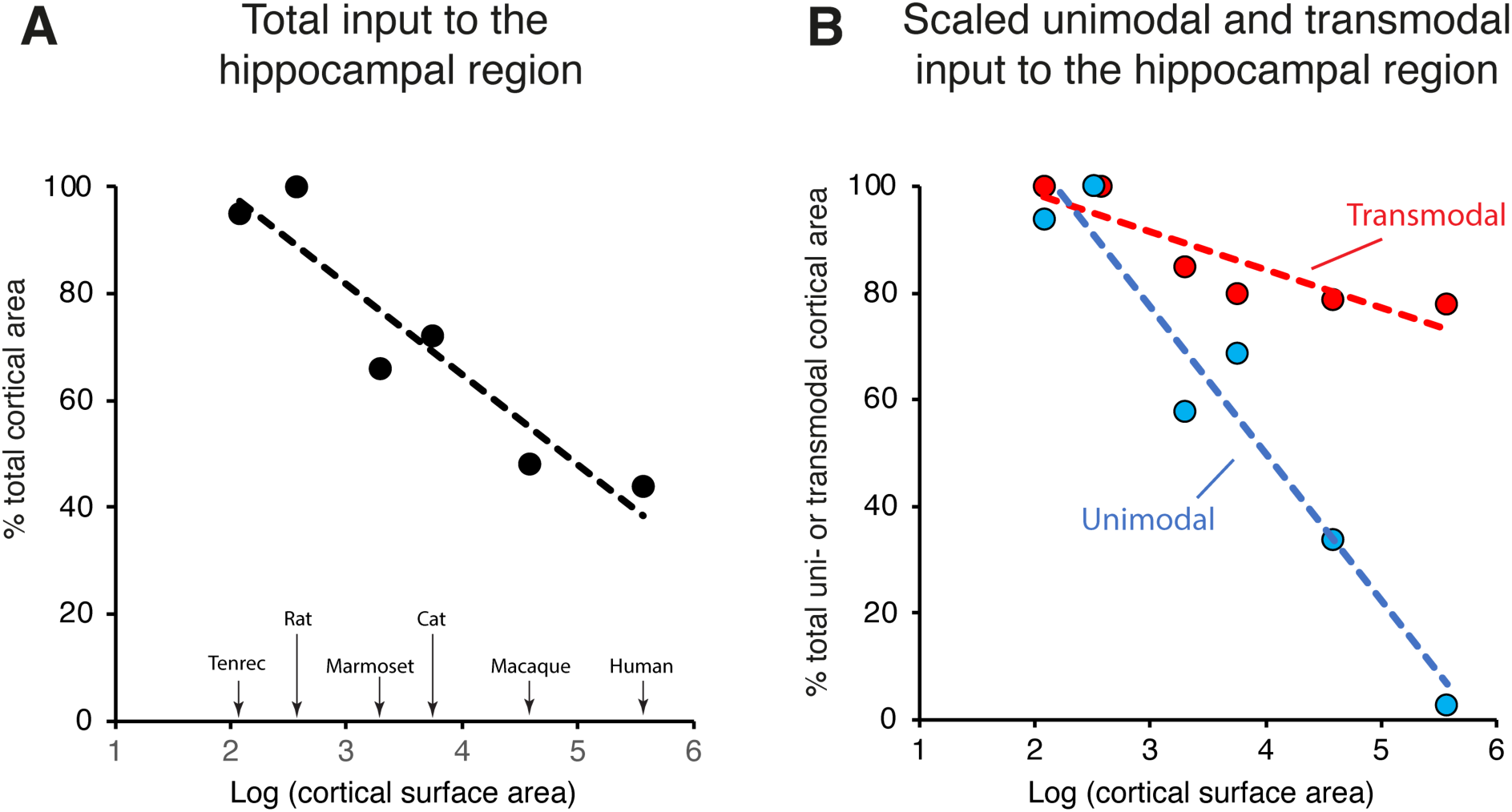
Cortical input to the hippocampal region. **A.** We find that the percentage of total cortical area (collapsed across unimodal and transmodal cortices) that projects to the hippocampal region decreases across species with the increase in brain size (Pearson correlation coefficient r=-0.92, p<0.01). In small brain mammals (the tenrec and rat), almost the entire cortical mantle projects directly to the hippocampal region. In primates, about half of the cortical mantle projects directly to the hippocampal region. **B.** Percentage of cortical input to the hippocampal region calculated separately from the total unimodal and transmodal cortical areas in each species. This analysis demonstrates that even though there is a general decrease in the percentage of total cortical area projecting to the hippocampal region, this decrease disproportionally targets unimodal input, while input from transmodal cortical areas remains relatively preserved with the increase of brain size (repeated measures ANCOVA, p<0.01).

Intriguingly, when we examined the reduction of input from unimodal sensory regions separately for each sensory modality, two phenomena were observed. First, direct anatomical input from sensory regions was eliminated gradually across species. Second, elimination of anatomical input from primary sensory regions was followed by elimination of anatomical input from non-primary sensory regions. Namely, in the rat, all primary and non-primary sensory regions project to the hippocampal region. However, in the cat, projections from primary somatosensory regions are not present, while projections from non-primary somatosensory, as well as from primary and non-primary auditory and visual regions remain. In the macaque, projections from primary auditory and visual regions (which are present in the cat) are absent, leaving only projections from non-primary auditory (caudal STG), non-primary visual (areas V4, IT cortex) and non-primary somatosensory (SII, granular insula) regions. The process of gradual elimination of anatomical connectivity with sensory regions culminates in humans, where connections with non-primary sensory regions that are present in the macaque are further eliminated. One notable exception to this pattern are connections of the hippocampal region with the piriform cortex (see Discussion).

## Discussion

In the current study we explored the evolutionary principles governing changes in anatomical connectivity between the cerebral cortex and the hippocampal region. To this end, we examined anatomical data from six mammalian species in which anatomical connections of the hippocampal regions are known. Our results point to a global reduction in cortical input to the hippocampal region with the increase in brain size. Critically, this reduction selectively targeted anatomical input from unimodal sensory regions, while input from transmodal cortical regions was relatively preserved. Furthermore, we observed that the elimination of unimodal sensory input was highly structured, such that direct input from primary sensory regions was eliminated first, followed by elimination of direct input from non-primary sensory regions.

Previous cross-species examinations of anatomical connectivity demonstrated that direct connections between neurons throughout the cortex are reduced with the increase in brain size, leading to the progressive segregation of the cortical circuitry into subnetworks (Herculano-Houzel et al., 2010; Palmer & Rosa, 2006; Ringo, 1991; M. G. P. Rosa et al., 2019). This progressive disconnection potentially reflects the necessity to balance between the number of neurons and efficient information processing, as well as intrinsic limitations dictated by available membrane space for synapses (Luppi et al., 2024; Stevens, 1989). The overall reduction in neural connectivity may be associated with the necessity to regulate the total amount of energy consumed by the brain, where most of the consumed energy is used for synaptic transmission (Harris et al., 2012). Additionally, most long-range anatomical projections in studied mammals are myelinated (Liewald et al., 2014), which imposes additional connectivity-related high energetic costs necessary for myelin synthesis and oligodendrocytes maintenance (Harris & Attwell, 2012; Saab et al., 2013). However, our results indicate that at least when connectivity with the hippocampal region is considered, the global reduction in neural connectivity selectively targets connections with primary and non-primary sensory regions. Specifically, our results show that the changes in cortical input to the hippocampal region reflect a continuous transition - from nearly all primary and non-primary sensory regions that project to the hippocampal region in small-brained mammals - the tenrec and rat - through only non-primary auditory, visual and somatosensory areas that project to the hippocampal region in the macaque, culminating in humans, which show no connectivity with any of the non-primary sensory regions.

While we currently do not have an answer to why specifically connections with unimodal sensory regions are eliminated, the observed trend is dramatic and indicates that hippocampus operates on fundamentally different types of information in different species. Since at least some of the connections between the hippocampal region and the parietal, retrosplenial, cingulate, and limbic cortices observed in the human exist already in small-brained mammals, we suggest that the gradual elimination of unimodal input through mammalian evolution reflects selective preservation of anatomical connectivity favouring input from transmodal regions.

One notable exception to the elimination of direct sensory input to the hippocampal region is input from the piriform cortex. Unlike other sensory modalities that were gradually disconnected from the hippocampal region, piriform cortex directly projects to the entorhinal cortex in the rat, cat and macaque, despite dramatic differences in the brain size and cortical complexity across species. While there is no evidence showing direct projections from the piriform cortex to the hippocampal region in the tenrec (personal correspondence with Heinz Künzle), it is plausible that these connections exist based on anatomical data from other species. In humans, the piriform cortex is typically assigned to the limbic network (Thomas Yeo et al., 2011), which may be part of the canonical default network (Girn et al., 2024). Therefore, such connections may exist in humans as well (Angeli et al., 2023; Reznik et al., 2023, 2024; Zheng et al., 2021). One possible explanation to the preserved connectivity with the piriform cortex is that unlike other primary sensory cortices, which do not share a direct anatomical boundary with the hippocampal region, the piriform cortex is anatomically adjacent to and developmentally associated with the perirhinal and entorhinal cortex. Taken to the extreme, in the tenrec, the anatomical boundary between the piriform cortex and the entorhinal cortex is not clearly defined (Künzle & Radtke-Schuller, 2000). With the dramatic increase in brain size across species, anatomical proximity of the piriform cortex barely changed, and it remained positioned closely to the perirhinal and entorhinal cortex, thus potentially retaining its anatomical connectivity with the hippocampal region. It is important to note that whereas in rats olfactory input innervates the full extent of the entorhinal cortex, in the macaque only about 15% percent of the entorhinal cortex is the recipient of direct olfactory input (Insausti et al., 2002).

It is intriguing to consider potential functional implications of such differences in cortical input to the hippocampal region across species. In the early levels of cortical processing, primary and non-primary sensory cortices encode different features of sensory information originating in the extrapersonal environment. In contrast, at later levels of cortical processing, transmodal areas converge these low-level, modality-specific signals and form cross-modal associative representations. For example, the ventral stream of visual processing, beginning in primary visual cortex, transforms low-level visual features into increasingly complex transmodal representations in anterior temporal cortex (Mishkin & Ungerleider, 1982). Therefore, transmodal areas that are associated with the hippocampal region convey to the hippocampal region highly processed multisensory information which is not tuned to a single stimulus property, such as the size of the respective representation on the retina or sound frequency. Our findings imply that the synaptic distance between sensory perception and mnemonic encoding dramatically differs across species, which could be reflected in cross-species differences in memory-related behaviours. The hippocampal region in the broad sense is known to be involved in episodic memory, learning and navigation (Eichenbaum, 2017; Igarashi et al., 2014; O’keefe & Lynn Nadel, 1978). These seemingly distinct functional properties of the hippocampus are typically considered separately and species-specific. However, they can also be seen in tandem, representing a functional continuum of a hierarchical increase in mnemonic complexity across species (Eichenbaum, 2017; Jeffery et al., 2024). For example, early memory theories suggested an existence of multiple memory systems that can be classified, for example, into “associative memory”, “representational memory” and “abstract memory” (Oakley, 1981). Importantly, these systems were proposed to be phylogenetically dependent on each other, such that different systems emerged at different stages of animal evolution (Tulving, 1985). More recent accounts of spatial navigation suggest a common framework that can account for the rich behavioral and experimental variability across different species (Jeffery et al., 2024). According to this framework, human memory can be studied as a form of navigation (albeit in abstract spaces (Bellmund et al., 2018), even though these two allegedly distinct functions are typically tested by different experimental paradigms.

Our current results suggest that the notion of memory may have changed through evolution from supporting sensory-based functions, such as response inhibition (O’keefe & Lynn Nadel, 1978), to supporting sensory-detached functions, such as episodic construction, future thinking and mental time travel (Buckner & Carroll, 2007; Hassabis & Maguire, 2007). It is difficult to imagine mnemonic processing driven by low-level sensory features (e.g., in the tenrec or rat); nevertheless, animal studies show that marmoset monkeys perform better in visual memory tasks requiring object-centred perception compared with rats, which are more prone to viewer-centred bias (Kell et al., 2023). Even though it is challenging to compare visual perception across species due to dramatic differences in the organization of the visual system (e.g., rats have a very wide field of view and lack a fovea (De Araujo et al., 2001; Solomon & Rosa, 2014)), this finding is consistent with reduced visual-mnemonic hierarchy in the rat compared with the marmoset (X.- J. Wang et al., 2020), suggesting that in rats, mnemonic fidelity to low-level sensory features potentially hinders object-centred generalization. We postulate that the transition from sensory-based cognition to increasingly abstract cognition (Tolman, 1948) is associated with the increasingly dominating role of transmodal input to the hippocampal region. Intriguingly, the distributed circuitry between transmodal cortical regions and its detachment from sensory processing was previously proposed to be especially expanded in humans and suited especially well for “internal mentation” functions, with some of them being highly dependent of the hippocampus, such as episodic recall, imagery and future thinking (Buckner & Krienen, 2013). Our current findings outline the potential evolutionary trajectory underlying the development of this circuitry into the human form (Figure 3).

**Figure 3.**
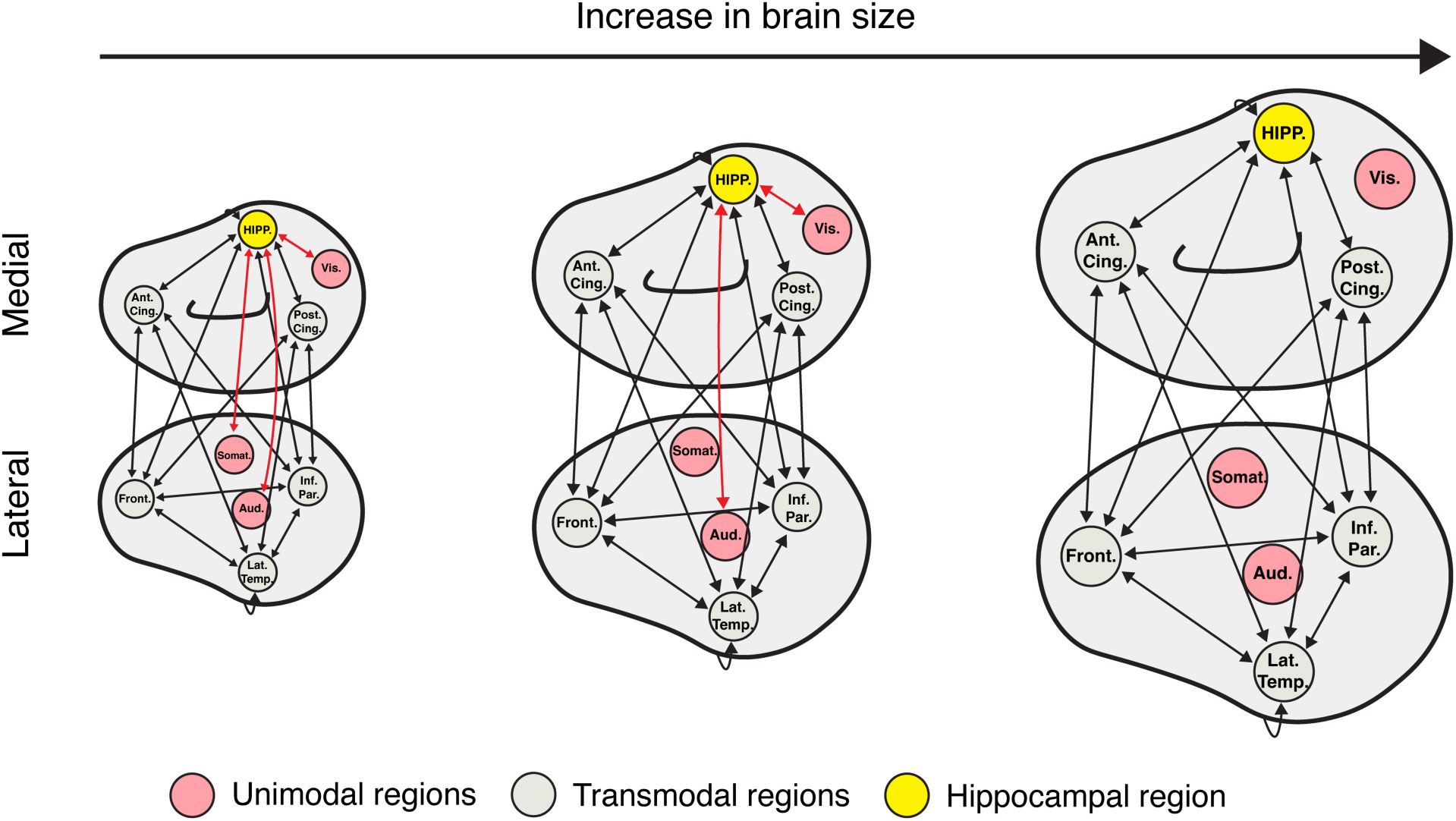
A schematic model describing the evolutionary trajectory of hippocampal circuitry. In small brain mammals, nearly all cortical regions project to the hippocampal region (denoted in the figure as HIPP). With the increase in brain size, direct projections from unimodal regions (red) gradually disappear, but direct projections from transmodal regions (grey) remain. When the cortical size reaches the level of human cortex, no connections between the hippocampal region and unimodal regions exist, and the hippocampal circuitry is almost exclusively associated with transmodal cortical regions. Ant. Cing. – Anterior cingulate; Aud. – Auditory cortex; Post. Cing. – Posterior cingulate cortex; Lat. Temp. – Lateral temporal cortex; Somat. – Somatosensory cortex; Inf. Par. – Inferior parietal cortex; Vis. – Visual cortex.

It is important to consider three main limitations pertaining to the current study. The first limitation is our sample size. We analyzed data from only six species - the only species with known distributed connectivity of the hippocampal region - four species representing Euarchontoglires (the rat, marmoset, macaque, and human), one species representing Afrotheria (the tenrec) and one species representing Laurasiatheria (the cat). This narrow sample size limited our ability to examine broad cross-taxa associations. Therefore, we are cautious in claiming that the observed results represent a general principle in mammalian phylogeny. For example, certain mammals, such as cetaceans, show a remarkably underdeveloped hippocampal region (Patzke et al., 2015), suggesting that at least some mammalian species might use other mechanisms for memory-related tasks. The second limitation is the indirect estimation of anatomical connectivity in humans using fMRI connectivity methods. Even though these methods were proved to be valid tools for non-invasively studying neuroanatomical connectivity (Vincent et al., 2007), unlike tract-tracing methods in the tenrec, cat, rat, marmoset and macaque, they are a proxy for measuring neuroanatomical connections. Therefore, absence of functional connectivity between brain regions in humans cannot indicate lack of anatomical connectivity (see for example (Reznik et al., 2023), for direct testing of connections between early sensory regions and the human hippocampal region). Nevertheless, a similar limitation holds also for the animal tracing experiments which often contain ‘‘hidden’’ data, either unpublished or not looked for. Therefore, no evidence for anatomical connectivity between certain regions in animal studies does not mean that this connectivity does not exist. The third limitation of our study is the definition of unimodal and transmodal cortical areas, which is particularly challenging given evidence pointing to multisensory properties of early sensory cortices (Schroeder & Foxe, 2005). However, since the functional properties of low-level multisensory integration are not yet clarified and their anatomical and physiological profiles contrast with those in “classic” multisensory areas (e.g., ventral intraparietal sulcus and superior temporal sulcus) (Smiley et al., 2007), in the current study we treated early sensory cortices as unimodal. Moreover, we adopted a neuroanatomical definition of unimodal sensory regions based on oligosynaptic access to information channelled by the sensory organs (e.g. retina, cochlea, tactile receptors).

To conclude, our study provides preliminary evidence for a continuous trend towards an increased role of transmodal input in mnemonic processing during 100 million years of mammalian evolution culminating in humans. Our findings suggest that the hippocampal region operates on fundamentally different types of input in different species, which would lead to specific hypotheses addressing differences in mnemonic representations in different species. Finally, our study suggests that it is critical to adopt an evolutionary perspective into the study of the hippocampal region, especially when cross-species data are considered (Jeffery et al., 2024; Martinez-Trujillo, 2025; Quian Quiroga, 2023; Zhu et al., 2023).

## Author contributions

D.R.: conceptualization, formal analysis, writing – original draft preparation, reviewing and editing; P.M.: resources, writing – reviewing and editing; M.G.P.R.: resources, writing – reviewing and editing; M.P.W.: writing – reviewing and editing; C.F.D.: funding acquisition, writing – reviewing and editing. All authors read and approved the final version of the manuscript.

## Acknowledgments

The authors thank Heinz Künzle, Casper Kerrén, Matan Mazor and Andrej Bicanski for helpful comments and discussions. This work is supported by the Max Planck Society. C.F.D. and M.P.W are supported by the Kavli Foundation. M.G.P.R. is supported by the Australian Research Council (DP210101042, DP210103865) and National Health and Medical Research Council (APP1194206). P.M. is supported by the Polish National Science Centre (2019/35/D/NZ4/03031). D.R is supported by the Kavli Foundation and by the European Research Council (grant agreement: 101220569).

