## Supplemental data for "Phylogeny of cortical-hippocampal projections reveals selective elimination of input from sensory regions"

Supplementary Information

<u>Cortical specialization across species</u>

We mapped cortical areas dedicated to unimodal and transmodal processing in six species – the tenrec, rat, cat, marmoset, macaque and human. Unimodal cortical areas were defined anatomically, as having the shortest connection path from sensory thalamic nuclei (with the exception of olfaction) and correspond to areas that are known to predominantly respond only to one type of modality, even if these responses are sometimes modulated by other senses. In each species, unimodal areas included visual, auditory, olfactory and somatosensory cortices. Transmodal areas included cortical areas that do not to show specificity to any single modality, which included polymodal and limbic cortices. The unimodal and transmodal cortical areas used for the analysis are presented in Tables S1-S2.

**Table S1.** Unimodal and transmodal cortical areas in the tenrec, rat and cat.

| **Unimodal** | **Transmodal** |
|---|---|
| **Tenrec** | |
| Piriform cortex | Cingulate area |
| Visual areas (V1 and rostral visual area) | Retrosplenial area |
| Somatosensory area (SI and SII) | Frontal area |
| Auditory area | |
| **Rat** | |
| Somatosensory cortex (primary, secondary) | Orbital cortex |
| Visual cortex (primary, secondary) | Retrosplenial cortex |

| | |
|---|---|
| Auditory cortex (primary, secondary) | Limbic cortex |
| Visceral cortex | Ventral temporal association cortex |
| Gustatory cortex | Posterior parietal association cortex |
| Piriform cortex | Agranular insular cortex |
| Periamygdaloid cortex | |
| **Cat** | |
| Visual areas (17, 18, 19, 21, AEV, ALLS, ALMS, CVA, DLS, PLLS, PLMS, PS, SVA, VLS) | Prefrontal cortex |
| Somatosensory areas (1, 2, 3, SII, SIII, SIV, SV) | Area 7, AES |
| Auditory areas (AI, AII, AAF, PE, DZ, FAES, auditory insula, PAF, TE*, VAF) | Lateral area 5, rest of area 5 |
| Lateral area 6 | Medial area 6 |
| Gustatory cortex, pyriform cortex | Cingulate, prelimbic and retrosplenial areas |

For parcelling the tenrec brain, we adopted prior work, which identified potential primary and secondary sensory areas (Krubitzer et al., 1997), as well as possible retrosplenial, cingulate and frontal areas (Künzle, 1995; Künzle & Rehkämper, 1992). In the cat, we followed the work by Payne et al. (Payne, 1993) and defined cat area 20 as the homologue of the primate parahippocampal cortex (postrhinal in the rodent).

**Table S2.** Unimodal and transmodal cortical areas in the marmoset, macaque and human.

| Unimodal | Transmodal |
|---|---|
| **Marmoset** | |
| Visual occipital areas (V1, V2, V3, V3A, V4, V4t, MT, prostriate cortex) | Prefrontal cortex (areas 8, 9, 10, 11, 12, 13, 14, 45, 46, OPro, OPAI, FEF) |
| Visual parietal areas (MST, V6, V6A, A19DI, A19M) | Parietal cortex (LIP, VIP, MIP, PGM, PG, PEC, Opt) |
| Visual temporal areas (FST, IT) | STP, TPpro, PGa/IPa |
| Somatosensory cortex (areas 1, 2, 3, PE, SII, granular insular cortex) | PF, PFG |
| Auditory (A1, caudal STG) | Ventral area 6 |
| Pyriform cortex | Auditory belt areas, STR |
| Periamygdaloid cortex | Limbic cortex (23, 24, 25, 29, 30, 32) |
| Gustatory cortex | Agranular and dysgranular insular cortex |
| **Macaque** | |
| Visual occipital areas (V1, V2, VP, V3A, V4, VOT, V4t, MT, prostriate cortex) | Prefrontal cortex (areas 9, 10, 11, 12, 13, 14, 45, 46, Pro, Pall, FEF) |
| Visual parietal areas (MST, PO, PIP, DP) | Parietal cortex (AIP, LIP, VIP, MDP, 7a) |
| Visual temporal areas (FST, IT) | STP, temporal pole |
| Somatosensory cortex (areas 1, 2, 3, 5, SII, granular insular cortex) | Area 7b |
| Auditory (A1, caudal STG) | Ventral area 6 |

| | |
|---|---|
| Pyriform cortex | Auditory belt areas and rostral STG |
| Periamygdaloid cortex | Limbic cortex (23, 24, 25, 29, 30, 32) |
| Gustatory cortex | Agranular and dysgranular insular cortex |
| **Human** | |
| Visual networks | Salience network |
| Somatomotor networks | Frontoparietal network A, B and C |
| Auditory network | Default networks |
| Premotor-Posterior Parietal Rostral network | Language network |
| Dorsal Attention Network A | Dorsal Attention Network B |
| Piriform cortex (assumed to be part of the limbic network) | Limbic network (Thomas Yeo et al., 2011) or the "grey" network (Du et al., 2024) |
| | Cingulo-Opercular network |

Due to functional ambiguity, marmoset cortical areas TPro, TF and TFO remained unclassified. Human data were based on the 15-network parcellation (Du et al., 2024); note that the human piriform cortex is believed to occupy the cortical area at the border of the temporal and frontal lobes, around the temporal pole (Ding et al., 2009) and is likely part of the cortical network previously referred to as the limbic network (Thomas Yeo et al., 2011). Assignment of the Dorsal Attention Network A and Dorsal Attention Network B to unimodal and transmodal processing was based on connectivity data examining whole-brain distributed topography associated with subregions of the visual system and the human medial temporal lobe (Reznik et al., 2023).

Relative size of unimodal and transmodal regions across species.

Percentage of each cortical modality in the tenrec was calculated based on electrophysiological data examining sensory response patterns in tenrec neocortex (Krubitzer et al., 1997). To this end, we counted the average number (across three presented animals) of electrodes placed in tenrec neocortex that showed either unimodal (auditory, visual or tactile) or transmodal (a combination of two or more sensory modalities, or no response at all) response patterns. The average number of electrodes for each response pattern are presented in Table S3.

**Table S3.** Number of electrodes that showed different response patterns in tenrec neocortex.

| Posterior visual | Anterior visual | Auditory | Somatosensory | Transmodal |
|---|---|---|---|---|
| 16.67 | 13 | 4.33 | 29.33 | 21.67 |

Percentage of each cortical modality in the rat was calculated based on a previously reported surface area estimation of different cortical subdivisions (see Table 3 in (Burwell et al., 1995)).

Percentage of each cortical modality in the cat was calculated based on the number of voxels assigned to each cortical subdivision in an MRI parcellation atlas of the cat brain (Stolzberg et al., 2017). See Table S4 for the size of each cortical subdivision in the cat, measured by the number of voxels.

**Table S4.** Number of voxels assigned to each cortical subdivision in the cat.

| Area | Size | Area | Size | Area | Size | Area | Size |
|---|---|---|---|---|---|---|---|
| A1 | 934 | 3b | 606 | CGa | 193 | 4sfu | 565 |
| A2 | 1486 | 5al | 439 | CGp | 1193 | 6aa | 541 |
| AAF | 1013 | 5am | 640 | 7m | 219 | 6ab | 712 |
| dPE | 582 | 5bl | 665 | 7p | 818 | 6ag | 69 |
| DZ | 267 | 5bm | 448 | AEV | 512 | 6iffu | 356 |
| pPE | 139 | 5m | 1054 | ALLS | 501 | PFdl | 505 |
| FAES | 781 | S2 | 548 | AMLS | 480 | PFdm | 754 |
| IN | 1481 | S2m | 721 | CVA | 1200 | PFv | 1062 |
| iPE | 1236 | S3 | 483 | DLS | 108 | AId | 1412 |
| PAF | 577 | S4 | 848 | PLLS | 921 | AIv | 1390 |
| TE | 1880 | S5 | 745 | PMLS | 809 | PL | 1405 |
| VAF | 338 | 17 | 7627 | PS | 414 | G | 448 |
| vPAF | 189 | 18 | 1329 | SVA | 444 | Pp | 590 |
| vPE | 1422 | 19 | 3408 | VLS | 214 | RS | 735 |
| 1 | 1464 | 21a | 248 | 4d | 298 | | |
| 2 | 634 | 21b | 632 | 4fu | 1126 | | |
| 3a | 465 | 7a | 1200 | 4g | 110 | | |

Percentage of each cortical modality in the marmoset was calculated based on a marmoset brain template (Majka et al., 2016). Each cortical subdivision's volume was corrected for tissue shrinkage. The size (measured in $mm^3$) of each cortical subdivision in the marmoset if presented in Table S5.

**Table S5.** Size (in $mm^3$) of cortical subdivisions in the marmoset.

| Area | Size | Area | Size | Area | Size | Area | Size |
|---|---|---|---|---|---|---|---|
| A1/2 | 23.9 | A3b | 20.8 | AuML | 2.5 | Pir | 11.9 |
| A10 | 27.1 | A45 | 4.8 | AuR | 4.1 | ProM | 10.0 |
| A11 | 11.3 | A46D | 3.8 | AuRM | 1.9 | ProSt | 7.3 |
| A13L | 6.8 | A46V | 3.6 | AuRPB | 3.9 | ReI | 4.5 |
| A13M | 5.6 | A47L | 15.6 | AuRT | 3.6 | S2E | 8.5 |
| A13a | 1.1 | A47M | 5.2 | AuRTL | 2.0 | S2I | 5.7 |
| A13b | 2.5 | A47O | 5.2 | AuRTM | 2.1 | S2PR | 3.7 |
| A14C | 0.6 | A4ab | 33.9 | DI | 5.5 | S2PV | 4.8 |

| | | | | | | | |
|---|---|---|---|---|---|---|---|
| A14R | 4.8 | A4c | 2.3 | FST | 14.0 | STR | 10.1 |
| A19DI | 10.1 | A6DC | 12.8 | GI | 5.0 | TE1 | 14.6 |
| A19M | 10.9 | A6DR | 10.2 | Gu | 5.6 | TE2 | 19.2 |
| A23V | 7.6 | A6M | 10.9 | IPro | 5.9 | TE3 | 38.5 |
| A23a | 13.6 | A6Va | 11.1 | LIP | 14.1 | TEO | 25.0 |
| A23b | 14.9 | A6Vb | 3.6 | MIP | 8.5 | TPO | 19.4 |
| A23c | 4.9 | A8C | 3.5 | MST | 15.3 | TPPro | 14.9 |
| A24a | 5.8 | A8aD | 12.2 | OPAl | 6.0 | TPro | 1.6 |
| A24b | 7.5 | A8aV | 12.4 | OPro | 5.4 | TPt | 6.3 |
| A24c | 5.5 | A8b | 14.9 | OPt | 9.4 | V1 | 298.1 |
| A24d | 5.8 | A9 | 7.1 | PE | 27.9 | V2 | 131.2 |
| A25 | 5.1 | AI | 2.1 | PEC | 6.8 | V3 | 63.1 |
| A29a-c | 10.1 | AIP | 4.0 | PF | 5.3 | V3A | 7.5 |
| A29d | 7.4 | APir | 0.8 | PFG | 14.2 | V4 | 33.9 |
| A30 | 14.3 | AuA1 | 8.2 | PG | 11.2 | V4T | 13.9 |
| A31 | 9.1 | AuAL | 2.8 | PGM | 6.6 | V5 | 16.3 |
| A32 | 9.2 | AuCL | 2.5 | Pga/Ipa | 4.9 | V6 | 32.0 |
| A32V | 2.0 | AuCM | 7.5 | PaIL | 2.2 | V6A | 10.4 |
| A3a | 22.5 | AuCPB | 6.5 | PaIM | 3.8 | VIP | 2.6 |

Percentage of each cortical modality in the macaque was calculated based on a previously reported surface area estimations of each cortical subdivision (see Table 2 in (Felleman & Van Essen, 1991)).

Percentage of each cortical modality in the human was calculated based on the 15-network parcellation (Du et al., 2024) projected to a standard surface space. The size (measured by the number of surface vertices) of each cortical network in the human are presented in Table S6.

**Table S6.** Number of surface vertices assigned to each cortical network in the human.

| Network | Size | Network | Size | Network | Size | Network | Size |
|---|---|---|---|---|---|---|---|
| VIS-P | 2230 | AUD | 1371 | LANG | 2747 | VIS-C | 1157 |
| CG-OP | 3852 | PM-Ppr | 1746 | FPN-B | 1503 | SAL/PMN | 1807 |
| DN-B | 3550 | dANT-B | 2386 | FPN-A | 2413 | DN-A | 2419 |
| SMOT-B | 1848 | SMOT-A | 3550 | dANT-A | 2295 | NONE | 3486 |

Cortical projections to the hippocampal region across species

To estimate the proportion of cortical areas involved in unimodal and transmodal processing from the total cortical input to the hippocampal region in each species, we analyzed animal tract-tracing and human fMRI data examining connectivity of the entorhinal, perirhinal and hippocampal (postrhinal in the rodent) cortices with the broader cortex. Where possible, we considered projections from the unimodal and transmodal cortical areas to the hippocampal region; in species without direct data about cortical input to the hippocampal region, cortical projections were estimated indirectly (see below).

One major advantage of the marmoset tracing data is that it allowed us to examine the connections between the hippocampal region and almost all sensory systems – somatosensory, motor, visual, and auditory. Nevertheless, almost half of the marmoset cortical mantle was not covered by tracer injections. For example, there were no available injections into the piriform cortex and the orbital frontal cortex (except for area A11). To account for these cross-species differences in anatomical coverage, we inferred anatomical connectivity between the marmoset hippocampal region and the cortical areas *that were not injected with a tracer* if these regions were found to be anatomically connected in the macaque. These inferred connections included the piriform cortex, insular cortex, gustatory cortex, middle inferior temporal cortex (TE2 and TE3), fundus of superior temporal sulcus, A13, A14, A25, A30, A31, A45, A47 (medial and orbital parts), orbital proiso- and periallo-cortex, and temporopolar cortex.

Since no tract-tracing is feasible in humans, we used correlations in spontaneous brain activity patterns as a proxy to indirectly measure in vivo mono- and polysynaptic neuroanatomical connectivity. This method was proved to be a powerful tool in elucidating the anatomical organization of the brain both at the level of local circuitry (Kenet et al., 2003) and at the level of macro-scale network architecture (Vincent et al., 2007). Specifically, correlations in blood oxygenation level-dependent (BOLD) signal measured with functional magnetic resonance imaging (fMRI) during spontaneous brain activity (fixation task data, also known as "resting state") were found to mirror cortical pathways estimated using tract-tracing both in the macaque and marmoset. However, even though fMRI connectivity methods allow to indirectly measure anatomical organization at the whole-brain level, the unimodal connectivity of the human olfactory modality with the hippocampal region still remains unresolved. Since direct connections exist between the piriform cortex and the entorhinal cortex in the cat, rat and macaque, we assume this connectivity to be present also in humans.

Anatomical connectivity between the neocortex and the hippocampal region in the tenrec

Most of the tenrec cortex is dedicated to olfactory processing, with neocortex being extremely small (about 15mm$^2$; (Krubitzer et al., 1997). Despite the small neocortex, previous work identified putative primary and secondary sensory fields, as well as putative retrosplenial, cingulate and frontal fields. Importantly, putative perirhinal and entorhinal fields in the tenrec were identified as well. Due to the small size of neocortical areas in the tenrec and general ambiguity about the cortical organization of the tenrec neocortex, it is difficult to draw clear conclusions about connectivity between the distinct neocortical areas and the hippocampal region. This is especially true given the relatively large size of tracer injections in the tenrec neocortex, which potentially cover more than one cortical field. Nevertheless, all tracer injections, except for one injection (94-18W), into the anterior-posterior and medial-lateral extents of the tenrec neocortex resulted in labelling in the tenrec hippocampal region. Another injection (99-29W) did not result in connectivity patterns that were rated as "moderate" or above. However, an injection in a similar area in a different case (99-25W) showed at least "moderate" connectivity patterns (Künzle, 2003). One injection that did not result in any labelling in the parahippocampal region targeted the caudal-medial part of the tenrec neocortex, overlapping with the putative primary visual area. However, since injections overlapping with the rostral visual area (which is known to receive direct input from dorsal lateral geniculate nucleus (Künzle, 2009)), we concluded that tenrec parahippocampal region receives input from at least one primary visual area. Note that in the tenrec, there is limited data on thalamo-cortical connections (Künzle, 2009). Therefore, unimodal sensory fields in the tenrec were identified based on response patterns to sensory stimulation, connections with other brain regions, myelination patterns and relative anatomical location compared with other species (Krubitzer et al., 1997).

Input and output projections in the marmoset and human

In the current study we could directly examine projections from the broader neocortex to the hippocampal region only in the tenrec, cat, rat and macaque. In two other species, the marmoset and human, such projections were estimated indirectly. In the marmoset, anatomical tracing data included only retrograde injections covering about half of the cortical mantle, excluding the hippocampal region. Therefore, in this species we could examine only projections from the entorhinal, perirhinal and hippocampal cortices to the broader neocortex. Tracing data from the macaque suggest that most cortical projections to the hippocampal region are reciprocal with some of the reciprocating projections being weaker (Lavenex et al., 2002). Therefore, we inferred cortical projections to the marmoset hippocampal region based on the return projections from this region to the neocortex. Importantly, even though almost all of macaque cortical-to-hippocampal projections are reciprocal, many of the return projections are not. For example, while there are no known projections directly connecting the macaque primary visual cortex to the hippocampal region, projections from the parahippocampal areas TH/TF to the primary visual cortex do exist (Markov et al., 2014; Rockland & Van Hoesen, 1994). Another example comes from the auditory modality. While retrograde injections in the auditory caudomedial belt area resulted in labelled cells along the macaque entorhinal cortex (Smiley et al., 2007), retrograde injections directly into the entorhinal cortex resulted in no labelled cells in auditory areas (Insausti et al., 1987). These examples suggest that using output projections as an estimate of input projections might result in overestimation, but not underestimation, of total projections from the cortex to the hippocampal region (Theodoni et al., 2021). Importantly, if we remove the estimated cortical projections to the hippocampal region in the marmoset that do not exist in the macaque (visual cortices V1, V2, primary somatosensory cortex 3a, secondary somatosensory cortex, primary auditory cortex), the percentage of transmodal input in the marmoset will be similar to that of the macaque. Therefore, as a final estimate of marmoset projections from the neocortex to the hippocampal region, we averaged the percentage of input projections estimated solely on output projections and the percentage of input projections estimated using macaque anatomical priors. It is still an open question whether projections from early visual, somatosensory, and/or auditory cortices to the hippocampal region exist in the marmoset.

In the human, since anatomical tract-tracing is unfeasible, we used spontaneous intrinsic brain activity patterns as a proxy to indirectly measure in vivo mono- and polysynaptic neuroanatomical connectivity. Like we elaborated above, even though functional connectivity methods serve as a powerful means to noninvasively estimate large-scale anatomical connectivity, they cannot distinguish between the directionality of connections. Therefore, it is possible that the correlations between the human entorhinal cortex and the putative olfactory cortex in or around the temporal pole are output connections. Moreover, unlike in the macaque and the marmoset, precision imaging studies imply that human parahippocampal areas TH/TF are not associated with early visual cortex (Reznik et al., 2023) (but see (Bergmann et al., 2016)), potentially meaning that even projections from the hippocampal region to the primary visual cortex, which are present in the macaque and the marmoset, are absent in humans.

Marmoset connectivity data without macaque-based adjustment

To examine the effect of marmoset data adjustment, we performed the main analysis without averaging marmoset and macaque neocortical connectivity with the hippocampal regions. As presented in Figure S2, main results remained unchanged.

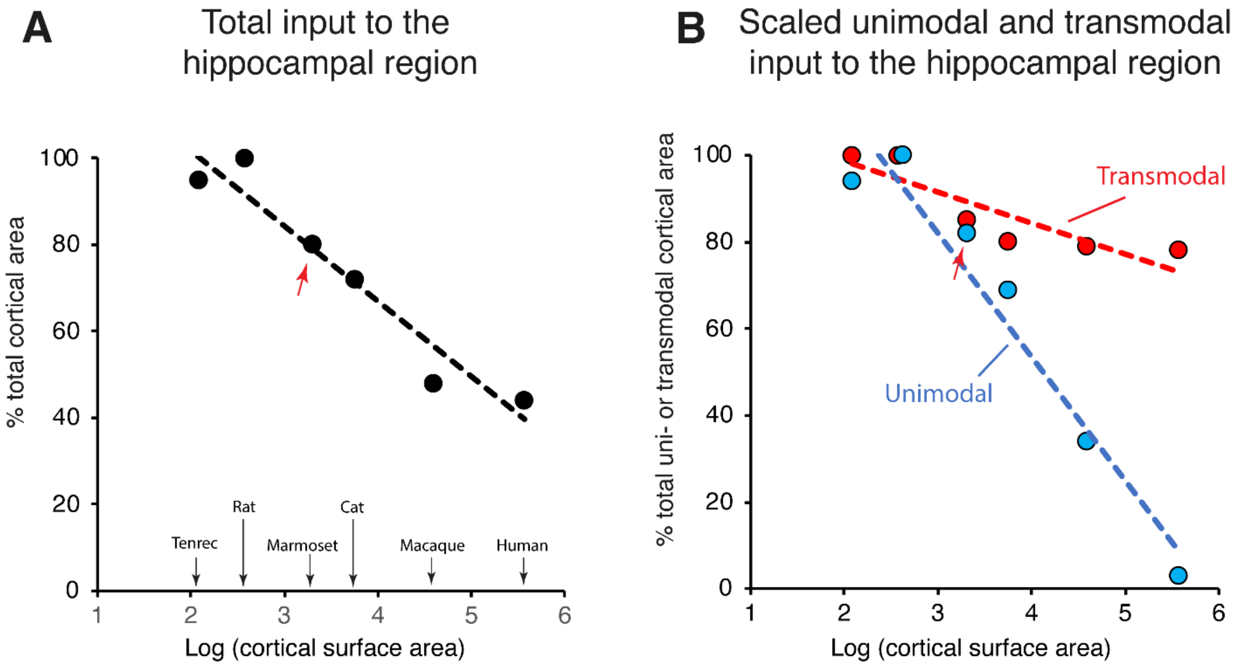


**Figure S1. Cortical input to the hippocampal region without adjusted marmoset data.**

**A.** We find that the percentage of total cortical area (collapsed across unimodal and transmodal cortices) that projects to the hippocampal region decreases across species with the increase in brain size (Pearson correlation coefficient $r=-0.96$, $p<0.01$). Changed marmoset data is marked with a red arrow.

**B.** Percentage of cortical input to the hippocampal region calculated separately from the total unimodal and transmodal cortical areas in each species. This analysis demonstrates that even though there is a general decrease in the percentage of total cortical area projecting to the hippocampal region, this decrease disproportionally targets unimodal input, while input from transmodal cortical areas remains relatively preserved with the increase of brain size (repeated measures ANCOVA, $p<0.05$). Changed marmoset data is marked with a red arrow.

Unimodal and transmodal input to the hippocampal region as a percentage of total cortical area

First, we examined differences in cortical specialization across species in our sample. Our results point to reduced unimodal and increased transmodal cortical areas percentage with the increase in brain size (Figure S1a; Pearson's r=-0.89, $p<0.05$). Note that this effect was driven by the difference in total unimodal/transmodal areas in the tenrec and human. Next, we examined direct anatomical projections to the hippocampal region separately for unimodal cortical regions and for transmodal cortical regions calculated as a percentage of total cortical area. We observed that total unimodal and transmodal input to the hippocampal region exhibited different patterns with the increase in species brain size. While total unimodal input reduced with the increase in brain size, total transmodal input increased with the increase in brain size (Figure S1b; repeated measures ANCOVA, $F_{(1,4)}=50.27$, $p<0.01$; note that the increase in transmodal input is driven by data from the tenrec and human).

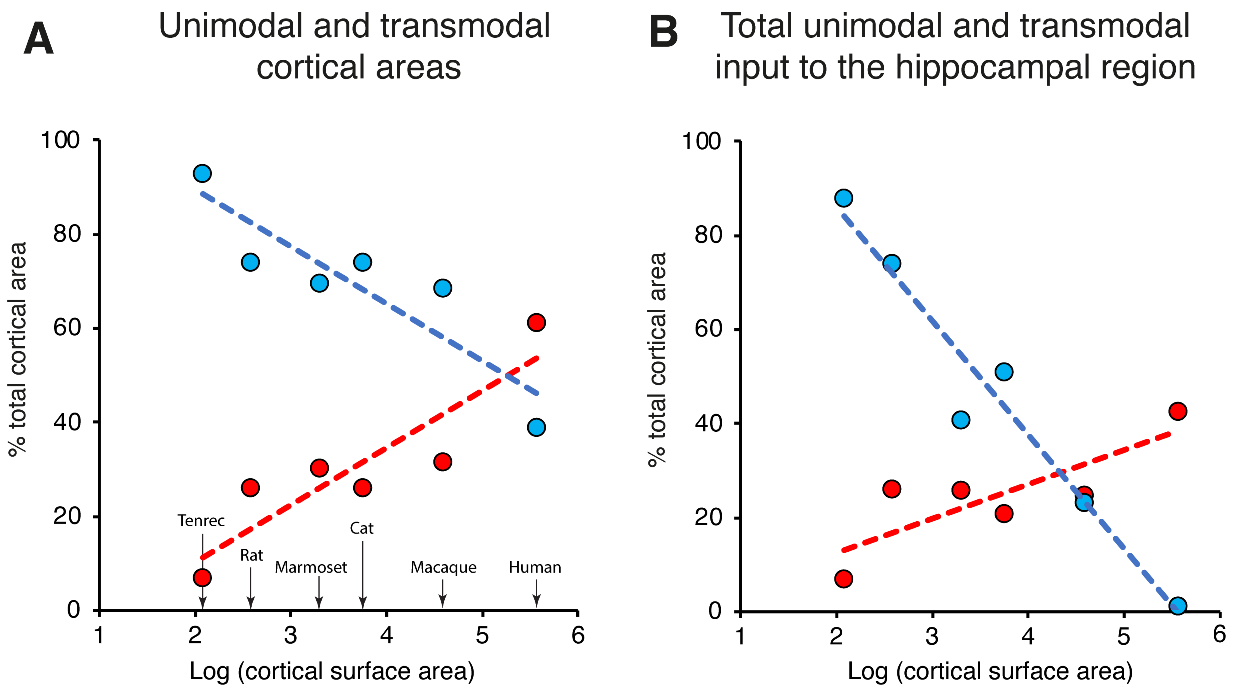

**Figure S2. Unimodal and transmodal areas and total cortical input to the hippocampal region.**

**A.** We find that the percentage of total cortical area involved in transmodal processing increased with the increase in species cortical surface area in our sample; correspondingly, the percentage of total cortical area involved in unimodal processing decreased across species (Pearson's $r=-0.89$, $p<0.05$). Note that this effect was driven by the difference in total unimodal/transmodal areas in the tenrec and human.

**B.** Cortical input to the hippocampal region calculated separately for unimodal and transmodal cortical areas as a percentage of total cortical area in each species point to different patterns with the increase in species brain size (repeated measures ANCOVA, $p<0.01$). Note that the increase in transmodal input to the hippocampal region across species is driven by tenrec and human data.

Cross-species analysis without human data

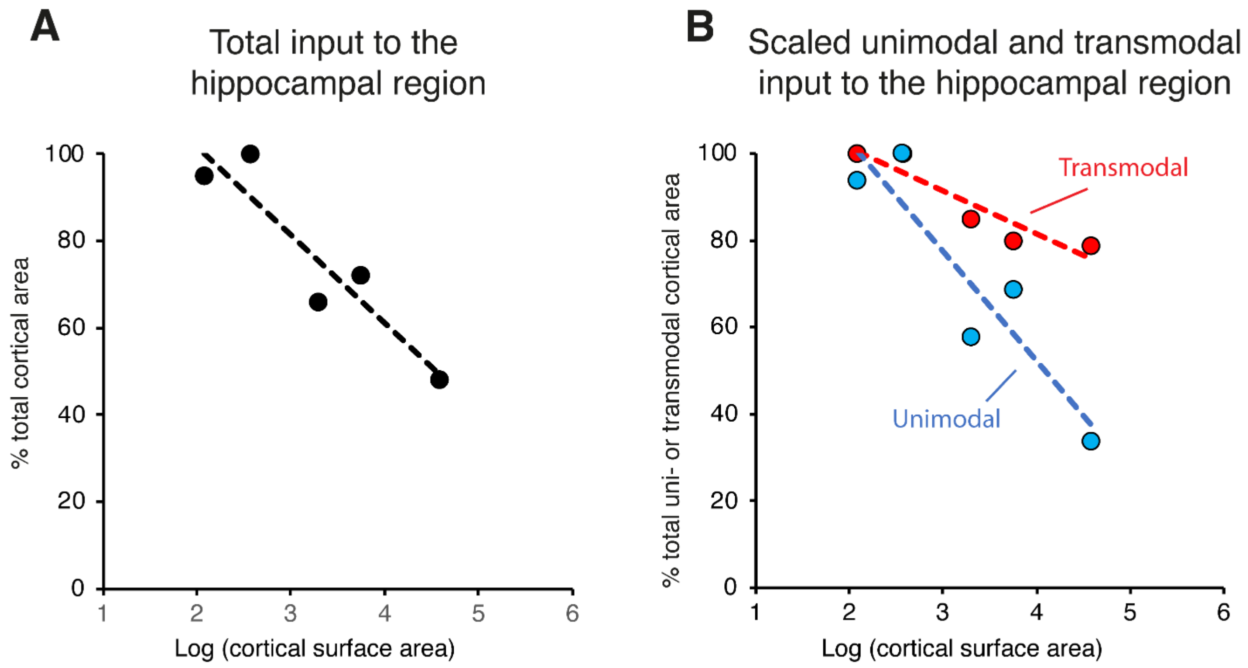


**Figure S3. Cortical input to the hippocampal region without human data**

**A.** After removing human data from the analysis, we still find that the percentage of total cortical area (collapsed across unimodal and transmodal cortices) that projects to the hippocampal region decreases across species with the increase in brain size (Pearson correlation coefficient $r=-0.93$, $p<0.05$). In small brain mammals (the tenrec and rat), almost the entire cortical mantle projects directly to the hippocampal region. In primates, about half of the cortical mantle projects directly to the hippocampal region.

**B.** Percentage of cortical input to the hippocampal region calculated separately from the total unimodal and transmodal cortical areas in each species. After removing human data from the analysis, differences in unimodal and transmodal input to the hippocampal region were no longer significant (repeated measures ANCOVA, $p=0.08$).

Bergmann, E., Zur, G., Bershadsky, G., & Kahn, I. (2016). The Organization of Mouse and Human Cortico-Hippocampal Networks Estimated by Intrinsic Functional Connectivity. *Cerebral Cortex*, *26*(12), 4497–4512. https://doi.org/10.1093/cercor/bhw327

Burwell, R. D., Witter, M. P., & Amaral, D. G. (1995). Perirhinal and postrhinal cortices of the rat: A review of the neuroanatomical literature and comparison with findings from the monkey brain. *Hippocampus*, *5*(5), 390–408. https://doi.org/10.1002/hipo.450050503

Ding, S. L., Van Hoesen, G. W., Cassell, M. D., & Poremba, A. (2009). Parcellation of human temporal polar cortex: A combined analysis of multiple cytoarchitectonic, chemoarchitectonic, and pathological markers. *Journal of Comparative Neurology*. https://doi.org/10.1002/cne.22053

Du, J., DiNicola, L. M., Angeli, P. A., Saadon-Grosman, N., Sun, W., Kaiser, S., Ladopoulou, J., Xue, A., Yeo, B. T. T., Eldaief, M. C., & Buckner, R. L. (2024). Organization of the human cerebral cortex estimated within individuals: networks, global topography, and function. *Journal of Neurophysiology*, *131*(6), 1014–1082. https://doi.org/10.1152/jn.00308.2023

Felleman, D. J., & Van Essen, D. C. (1991). Distributed Hierarchical Processing in the Primate Cerebral Cortex. *Cerebral Cortex*, *1*(1), 1–47. https://doi.org/10.1093/cercor/1.1.1

Insausti, R., Amaral, D. G., & Cowan, W. M. (1987). The entorhinal cortex of the monkey: II. Cortical afferents. *Journal of Comparative Neurology*, *264*(3), 356–395. https://doi.org/10.1002/cne.902640306

Kenet, T., Bibitchkov, D., Tsodyks, M., Grinvald, A., & Arieli, A. (2003). Spontaneously emerging cortical representations of visual attributes. *Nature*, *425*(6961), 954–956. https://doi.org/10.1038/nature02078

Krubitzer, L., K�nzle, H., & Kaas, J. (1997). Organization of sensory cortex in a Madagascan insectivore, the tenrec (Echinops telfairi). *The Journal of Comparative Neurology*, *379*(3), 399–414. https://doi.org/10.1002/(SICI)1096-9861(19970317)379:3<399::AID-CNE6>3.0.CO;2-Z

Künzle, H. (1995). Regional and Laminar Distribution of Cortical Neurons Projecting to Either Superior or Inferior Colliculus in the Hedgehog Tenrec. *Cerebral Cortex*, *5*(4), 338–352. https://doi.org/10.1093/cercor/5.4.338

Künzle, H. (2003). Neocortical connections with perihippocampal and periamygdalar regions in the hedgehog tenrec. *Anatomy and Embryology*, *207*(4–5), 389–407. https://doi.org/10.1007/s00429-003-0356-z

Künzle, H. (2009). Tracing thalamo-cortical connections in tenrecA further attempt to characterize poorly differentiated neocortical regions, particularly the motor cortex. *Brain Research*, *1253*, 35–47. https://doi.org/10.1016/j.brainres.2008.11.052

Künzle, H., & Rehkämper, G. (1992). Distribution of Cortical Neurons Projecting to Dorsal Column Nuclear Complex and Spinal Cord in the Hedgehog Tenrec, *Echinops telfairi*. *Somatosensory & Motor Research*, *9*(3), 185–197. https://doi.org/10.3109/08990229209144770

Lavenex, P., Suzuki, W. A., & Amaral, D. G. (2002). Perirhinal and parahippocampal cortices of the macaque monkey: Projections to the neocortex. *Journal of Comparative Neurology*, *447*(4), 394–420. https://doi.org/10.1002/cne.10243

Majka, P., Chaplin, T. A., Yu, H., Tolpygo, A., Mitra, P. P., Wójcik, D. K., & Rosa, M. G. P. (2016). Towards a comprehensive atlas of cortical connections in a primate brain: Mapping tracer

injection studies of the common marmoset into a reference digital template. *Journal of Comparative Neurology*, *524*(11), 2161–2181. https://doi.org/10.1002/cne.24023
Markov, N. T., Ercsey-Ravasz, M. M., Ribeiro Gomes, A. R., Lamy, C., Magrou, L., Vezoli, J., Misery, P., Falchier, A., Quilodran, R., Gariel, M. A., Sallet, J., Gamanut, R., Huissoud, C., Clavagnier, S., Giroud, P., Sappey-Marinier, D., Barone, P., Dehay, C., Toroczkai, Z., … Kennedy, H. (2014). A Weighted and Directed Interareal Connectivity Matrix for Macaque Cerebral Cortex. *Cerebral Cortex*, *24*(1), 17–36. https://doi.org/10.1093/cercor/bhs270
Payne, B. R. (1993). Evidence for Visual Cortical Area Homologs in Cat and Macaque Monkey. *Cerebral Cortex*, *3*(1), 1–25. https://doi.org/10.1093/cercor/3.1.1
Reznik, D., Trampel, R., Weiskopf, N., Witter, M. P., & Doeller, C. F. (2023). Dissociating distinct cortical networks associated with subregions of the human medial temporal lobe using precision neuroimaging. *Neuron*, *111*(17), 2756-2772.e7. https://doi.org/10.1016/j.neuron.2023.05.029
Rockland, K. S., & Van Hoesen, G. W. (1994). Direct Temporal-Occipital Feedback Connections to Striate Cortex (V1) in the Macaque Monkey. *Cerebral Cortex*, *4*(3), 300–313. https://doi.org/10.1093/cercor/4.3.300
Smiley, J. F., Hackett, T. A., Ulbert, I., Karmas, G., Lakatos, P., Javitt, D. C., & Schroeder, C. E. (2007). Multisensory convergence in auditory cortex, I. Cortical connections of the caudal superior temporal plane in macaque monkeys. *Journal of Comparative Neurology*, *502*(6), 894–923. https://doi.org/10.1002/cne.21325
Stolzberg, D., Wong, C., Butler, B. E., & Lomber, S. G. (2017). Catlas: An magnetic resonance imaging-based three-dimensional cortical atlas and tissue probability maps for the domestic cat ( *Felis catus* ). *Journal of Comparative Neurology*, *525*(15), 3190–3206. https://doi.org/10.1002/cne.24271
Theodoni, P., Majka, P., Reser, D. H., Wójcik, D. K., Rosa, M. G. P., & Wang, X.-J. (2021). Structural Attributes and Principles of the Neocortical Connectome in the Marmoset Monkey. *Cerebral Cortex*, *32*(1), 15–28. https://doi.org/10.1093/cercor/bhab191
Thomas Yeo, B. T., Krienen, F. M., Sepulcre, J., Sabuncu, M. R., Lashkari, D., Hollinshead, M., Roffman, J. L., Smoller, J. W., Zöllei, L., Polimeni, J. R., Fischl, B., Liu, H., & Buckner, R. L. (2011). The organization of the human cerebral cortex estimated by intrinsic functional connectivity. *Journal of Neurophysiology*, *106*(3), 1125–1165. https://doi.org/10.1152/jn.00338.2011
Vincent, J. L., Patel, G. H., Fox, M. D., Snyder, A. Z., Baker, J. T., Van Essen, D. C., Zempel, J. M., Snyder, L. H., Corbetta, M., & Raichle, M. E. (2007). Intrinsic functional architecture in the anaesthetized monkey brain. *Nature*, *447*(7140), 83–86. https://doi.org/10.1038/nature05758
Yan, X., Kong, R., Xue, A., Yang, Q., Orban, C., An, L., Holmes, A. J., Qian, X., Chen, J., Zuo, X.-N., Zhou, J. H., Fortier, M. V, Tan, A. P., Gluckman, P., Chong, Y. S., Meaney, M. J., Bzdok, D., Eickhoff, S. B., & Yeo, B. T. T. (2023). Homotopic local-global parcellation of the human cerebral cortex from resting-state functional connectivity. *NeuroImage*, *273*, 120010. https://doi.org/10.1016/j.neuroimage.2023.120010
